# Evidence for sex microchromosomes in a species with temperature-influenced sex determination (*Crotaphytus collaris*)

**DOI:** 10.1101/270983

**Authors:** Jodie M. Wiggins, Enrique Santoyo-Brito, Jon B. Scales, Stanley F. Fox

**Affiliations:** Department of Integrative Biology, Oklahoma State University, OK 74078, USA; Department of Biology, Midwestern State University, TX 76308, USA

**Keywords:** Sex-determining mechanism, genotypic sex determination, temperature-dependent sex determination, gene-dosage, sex microchromosome, reptile, *Crotaphytus collaris*

## Abstract

The characteristics of a species’ evolution can be powerfully influenced by its mode of sex determination and, indeed, sex determination mechanisms vary widely among eukaryotes. In non-avian reptiles, sex was long thought to be determined bimodally, either by temperature or genetics. Here we add to the growing evidence that sex determining mechanisms in reptiles fall along a continuum rather than existing as a mutually exclusive dichotomy. Using qPCR, we demonstrate that the lizard *Crotaphytus collaris* possesses sex-based gene dosage consistent with the presence of sex michrochromosomes, despite that extreme incubation temperatures can influence hatchling sex ratio. Our results suggest a temperature override that switches genotypic females to phenotypic males at high and low temperatures.

## Introduction

Mode of sex determination has far reaching consequences affecting evolutionary processes such as speciation (Haldane 1922), adaptive capability and heterozygosity (Bull 1983; Shine et al. 2002; Burt and Trivers 2006), maternal capacity for sex ratio manipulation (Kratochvíl et al. 2008), extent of secondary sexual characteristics (Reeve and Pfennig 2003) and capacity to respond to climate change (Mitchell and Janzen 2010). Among eukaryotes, the array of sex determination mechanisms (SDMs) is diverse (Bull 1983; Charlsworth 1996; Bachtrog et al. 2014). Environmental sex determination (ESD) is characterized by a mode of sex determination entirely dependent on environmental factors such as temperature encountered during embryogenesis (Merchant-Larios and Díaz-Hernández 2013), photoperiod (Walker 2005; Guler et al. 2012), or social cues during subsequent development (Bull 1983; Janzen and Paukstis 1991; Valenzuela and Lance 2004; Bachtrog et al. 2014). Conversely, sex in GSD species is determined at conception by genes, and is uninfluenced by environment (Bull 1983; Sarre et al. 2004).

Long held was the belief that within reptilian lineages there existed a single dichotomy of mutually exclusive SDMs (Bull 1983; Janzen and Paukstis 1991). Either a species’ sex was thought to be determined by the environment (environmental sex determination; ESD) or by sex chromosomes (genotypic sex determination; GSD) (Bull 1983; Janzen and Paukstis 1991; Bachtrog et al. 2014). In fact, reptilian species do utilize both male and female heterogamety (GSD) (King 1977) and temperature-dependent sex determination (TSD; a specific type of ESD) (Ewert and Nelson 1991). However, in recent years, the line between ESD and GSD has become decidedly blurred, and an increasingly complex picture is emerging in which GSD and TSD are the ends of a continuum of SDMs in reptiles. Examples of intermediate SDMs include species in which different populations utilize different sex determining mechanisms (Pen et al. 2010), temperature-dependent sex reversal of a species with a ZZ/ZW GSD system in the wild (Quinn et al. 2007; Holleley et al. 2015), and revelations of the genetic underpinnings of TSD in alligators and turtles (Spotila et al. 1998; Smith et al. 1999; Kettlewell et al. 2000; Western and Sinclair 2001). This shifting landscape provides an exceptional study system for better understanding sex determination in vertebrates (Sarre et al. 2004).

The collared lizard, *Crotaphytus collaris*, is a widespread, oviparous species in which sex chromosomes have not been identified (Gorman 1973; De Smet 1981). Yet, *C. collaris* has been classified as a GSD species based on phylogenetic analyses (Pokorna and Kratochvíl 2009). However, as mentioned above, even members of the same species can utilize different sex determining mechanisms (Pen et al. 2010). Thus, classifying *C. collaris* solely based on phylogeny may provide an incorrect or incomplete picture (Viets et al. 1994). Compellingly, in an investigation seeking to determine if *C. collaris* utilizes TSD, an inverse TSD type II pattern was identified in which the percentage of female offspring declined as constant incubation temperatures or treatments approached high and low extremes (Santoyo-Brito et al. 2017). While the authors of this study point out that their sample size was low, this indicates a temperature influence on sex determination in *C. collaris*. However, even at low and high treatments this study did not find ratios of either sex nearing 100%. These findings hint at a more complex mechanism for sex determination than pure TSD or pure GSD in *C. collaris*, as suggested in other reptile species (Quinn et al. 2007; Radder et al. 2008; Holleley et al. 2015). In species that possess XY sex chromosomes, the heterogametic sex is expected to have half the dosage of X-linked genes (Rovatsos et al. 2014a). Indeed, sex-specific gene dosage at two loci in the closely related *Crotaphytus insularis* points to heterogamety (males are heterogametic) and, thus, to the possibility of GSD in Crotaphytids (Rovatsos et al. 2014b). The apparent conflict between the findings of the above studies compels us to determine if *C. collaris* demonstrates gene dosage akin to that identified in *C. insularis*.

## Methods

### DNA isolation and PCR

Blood was collected from the toes of twenty wild caught lizards (10 male, 10 female) upon capture at Sooner Lake Dam, Pawnee Co., Oklahoma. Blood was immediately preserved on Whatman FTA classic cards. DNA was later extracted from the cards by excising a 3-mm square of blood-saturated card using sterile scissors then following the GE Healthcare extraction protocol using Chelex^®^ 100 resin. We tested for gene dosage in the genes ATP2A2 (sarcoplasmic reticulum calcium ATPase 2), TMEM (transmembrane protein 123D), and PEBP1 (phosphatidylethanolamine binding protein 1) (Table 1). Primer sequences for the genes TMEM, PEBP1, and ATP2A2 were obtained from Rosavatos et al (2014; Table 1). The single copy gene EF1*α* was used for gene dosage normalization. PCRs were assembled in 25-µl final volumes containing 12.5 µl 2X Bullseye EvaGreen qPCR master mix buffer (MidSci, St. Louis, MO), 25 ng genomic DNA, and 200 pMoles forward and reverse primers. The thermal profile was an initial denaturation of 10 min at 94°C followed by 40 cycles of 94°C for 30 sec, 55°C for 30 sec, 72° C for 45 sec. Amplification via qPCR was executed in a Stratagene MX3005 thermocycler.

**Table 1.**
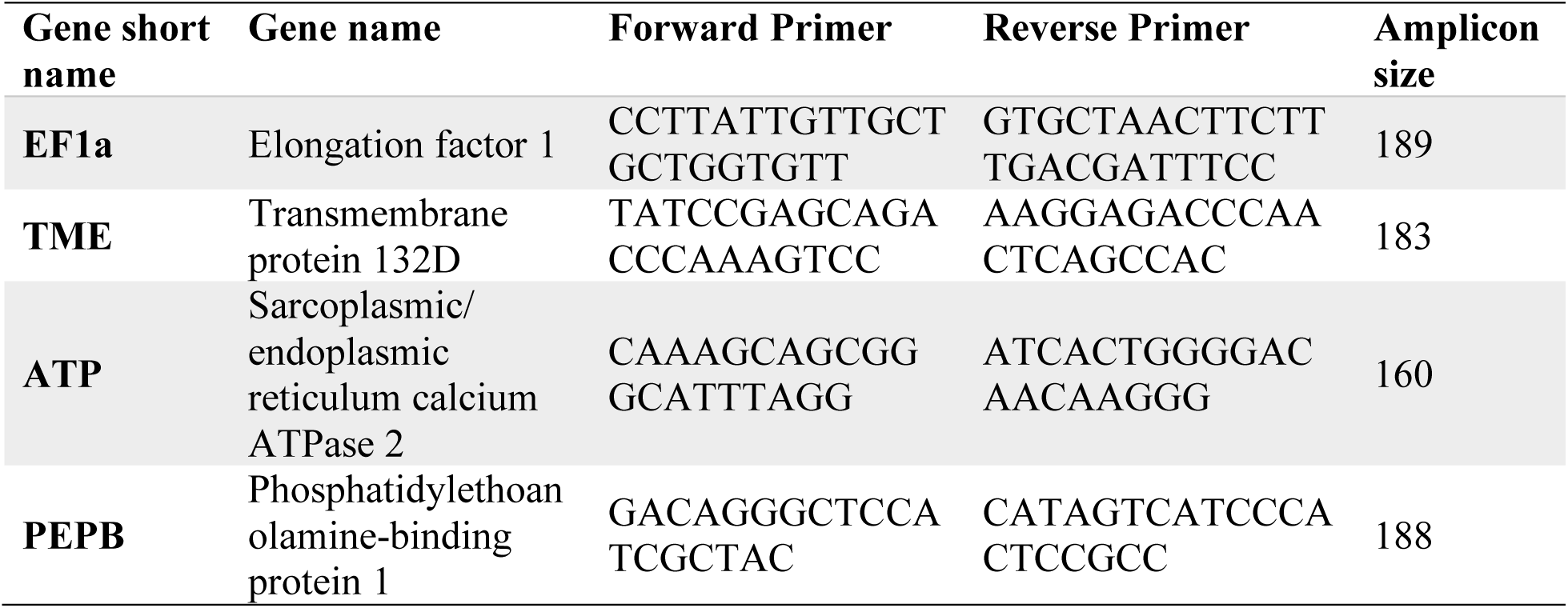
Genes and primer sequences used to determine relative gene dosage through qPCR in *Crotaphytus collaris*

### Gene Dosage Calculations

Final gene dosage ratios were calculated for the two male and two female lizards whose DNA consistently and reliably amplified across three replicates. Crossing point values were calculated in MaxPro (Stratagene) then normalized to EF1*α*. Gene dosage was calculated as in Rosavotas et al. (2014b) with: R = [2^Cp gene^/2^Cp EF1*α*^]^−1^ and r (relative gene dosage ratio) = R_male_/R_female_. It was expected that male *C. collaris* are the heterogametic sex based on results in the closely related *Crotaphytus insularis* (Rovatsos et al. 2014b), thus an r = 0.5 is expected for single copy, X-linked genes while r = 1.0 is expected for autosomal genes.

## Results

Results demonstrate sex-linked differences for all three analyzed genes (ATP2A2, TMEM, and PEBP1; Fig 1). In each case, the average female r value is exactly one. This result is consistent with females having two copies of the gene of interest and, thus, being the homogametic sex. For each gene an average value near 0.5 was obtained in the males (ATP2A2 = 0.59, TMEM = 0.61, PEBP1 = 0.59). This result is consistent with males having a single gene copy and being heterogametic.

**Figure 1.**
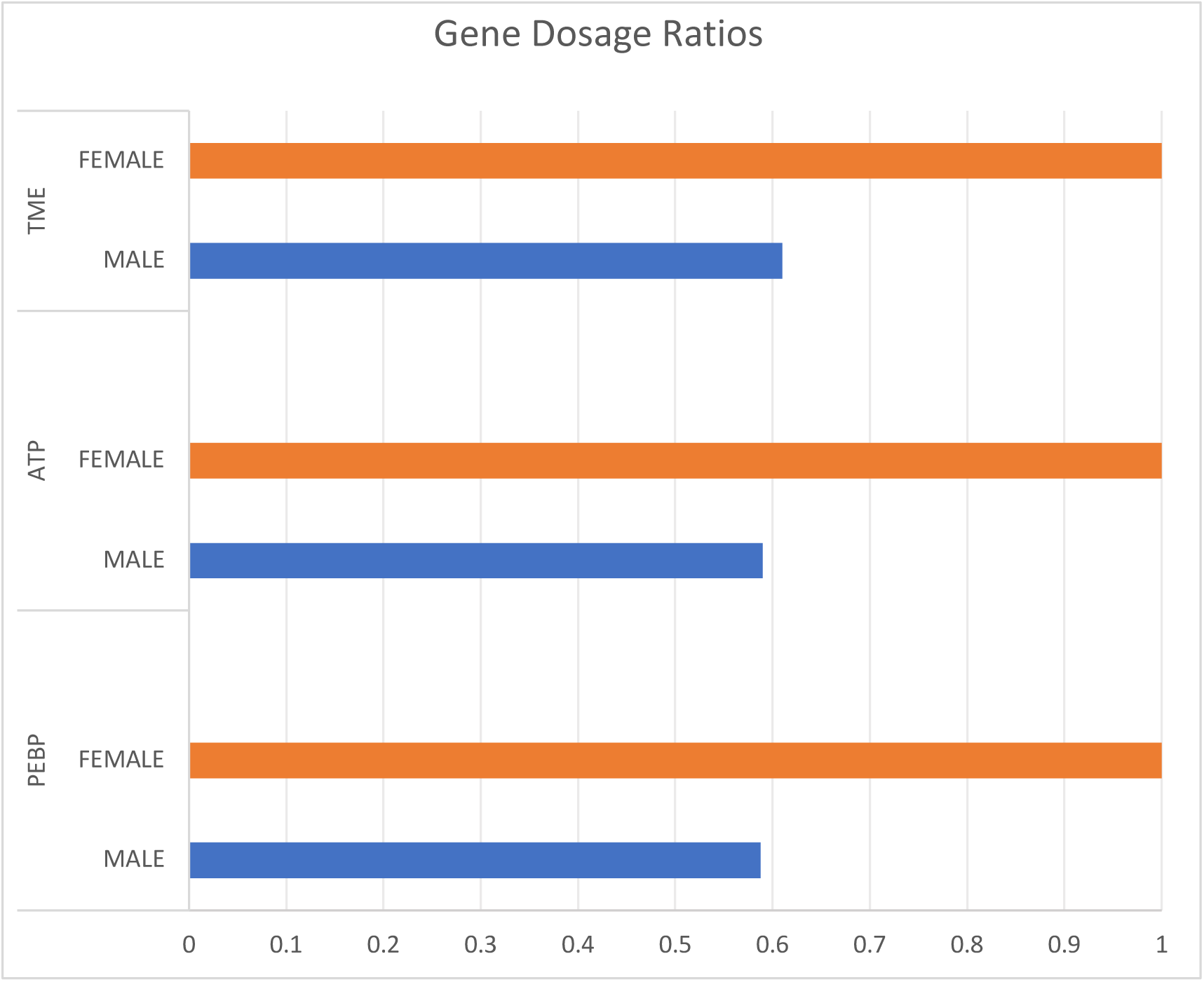
Relative gene dosage at genes PEBP, ATP, and TME for the two males and two females whose DNA amplified consistently and reliably across three replicates. For each gene, the female gene dosage is exactly 1 and the male gene dosage is near 0.5. Each of these genes maps to the *Anolis carolinensis* X microchromosome, thus, this pattern is consistent with male heterogamety.

## Discussion

Though sex chromosomes have not been detected in Crotaphytids (Gorman 1973; De Smet 1981) our results are consistent with heterogamety and point to the existence of GSD in *C. collaris* as in *C. insularis* (Rovatsos et al. 2014b). While GSD is common among lizards, *C. collaris* has been shown to experience changes in sex ratios with varying incubation temperatures (Santoyo-Brito et al. 2017). *Crotaphytus collaris* may be another in the emerging examples of a reptilian species with GSD that can be overridden by temperature extremes (Shine et al. 2002; Holleley et al. 2015). We agree with Santoyo-Brito et al. (2017) that *C. collaris* likely possess sex microchromosomes in an XX/XY pattern and that extreme incubation temperatures alter the sex determining pathway such that XX individuals develop as phenotypic males. We further hypothesize that gravid *C. collaris* females select nest sites such that GSD will function without temperature interference as evidenced by field hatchling ratios near 50/50 as expected in GSD populations (Wiggins unpublished data).

Evidence continues to emerge that some extant reptile species employ multimodal sex determining mechanisms (Shine et al. 2002; Valenzuela et al. 2003; Sarre et al. 2004; Quinn et al. 2007; Radder et al. 2008; Holleley et al. 2015). These species may be at a transition point from GSD to TSD. *Crotaphytus collaris* demonstrates gene dosage consistent with that of 28 species of Pleurodont iguanian lizards (Pleurodonta) spanning 11 genera and including *Anolis carolinensis* and *C. insularis* (Rovatsos et al. 2014a,b). The absence of genes on the Y that are present on the X, coupled with the chromosomal looping in the *A. carolinensis* XY synaptonemal complex, points to a degenerate Y microchromosome in this species and, subsequently, those with the same sex-based gene dosage (Alföldi et al. 2011; Bachtrog et al. 2014; Rovatsos et al. 2014a,b; Lisachov et al. 2017). Thus, there exists the strong possibility that *C. collaris* also possesses a degenerate Y microchromosome. Results from Santoyo-Brito et al. (2017) show a decline in number of females hatched at extreme high and low temperatures, pointing to a temperature override that converts XX individuals into phenotypic males (XXm). If these XXm are viable, they may mate and reproduce (as males). While this scenario will lead to an increase in the proportion of genetic females, the possibility of producing offspring who possess both degenerate chromosomes (YY) would be avoided as in *Bassiana duperreyi* (Shine et al. 2002). As a result, GSD and TSD could conceivably co-exist in *C. collaris* without a definitive transition to one or the other. However, as global temperatures rise, frequent shifts of XXf to XXm could eliminate the Y microchromosome, driving this species to pure TSD and increasing its vulnerability to extinction by fixation of homogamety and absence of XY individuals (Mitchell and Janzen 2010).

In summary, our data convincingly point to the presence of as-yet unidentified sex microchromosomes in *C. collaris* as suggested by Santoyo-Brito et al. (2017) and add to the growing evidence that SDMs in non-avian reptiles are not bimodal (Shine et al. 2002; Holleley et al. 2015). Further inquiry into the effects of extreme temperature incubations on *C. collaris* sex determination is warranted (Santoyo-Brito et al. 2017); specifically, investigating whether some of the individuals incubated at either high or low temperature extremes are genotypically female but phenotypically male as suggested by Santoyo-Brito et al. (2017) and as shown in free-ranging *P. vitticeps* (Holleley et al. 2015), if such individuals are reproductively viable, and if they exist in natural populations.

